# Proteome Microarray Screening Identifies Human Polyphosphate-Binding Proteins in the Phosphatidylinositol Signaling Pathway

**DOI:** 10.1101/2021.12.10.472157

**Authors:** Viola Krenzlin, Julian Roewe, Marcel Strueve, María Martínez-Negro, Christoph Reinhardt, Svenja Morsbach, Markus Bosmann

## Abstract

Polyphosphates are linear chains of orthophosphate residues that are present in all living cells. Polyphosphates are released from platelet δ-granules and are also produced in bacteria. Polyphosphates are procoagulant in mammalian species and in bacteria are required for energy and phosphate storage, stress resistance, chelation of metal ions and escaping host immunity. Despite these pleiotropic effects, sparse information is available on molecular binding partners of polyphosphates. Here, we used a slide-based human proteome microarray screen for the search of polyphosphate-binding proteins. This approach suggested several novel proteins with relation to the phosphatidylinositol signaling pathway. The highest signals were obtained for Disabled-1 (DAB1) and phosphatidylinositol-5-phosphate 4-kinase 2B (PIP4K2B). Isothermal titration calorimetry was used for confirmation of DAB1 interactions with long-chain polyphosphates. These results offer new rationale to further investigate the interference of polyphosphates with intracellular signaling pathways.

---

Inorganic polyphosphates are major modulators of blood coagulation *in vivo*.[1] Released from platelet δ-granules, these anionic polymers accelerate FXIIa-mediated FXI activation, thrombin generation, block TFPI activity, and strengthen fibrin clots by enhancing their mechanical stability and resistance to fibrinolysis.[2] In bacteria, polyphosphates are associated with energy and phosphate storage, stress resistance, chelation of metal ions and escaping host immunity.[3] Despite these emerging roles, the knowledge of polyphosphate-binding proteins remains incomplete, although such insights are crucial to better understand the pleitropic activities of polyphosphates in living cells.

Here, we report a screen using a human proteome microarray (HuProt™) to identify novel polyphosphate-binding proteins. The HuProt™ microarray (Cambridge Protein Arrays Ltd., Babraham Research Campus, Cambridge CB22 3AT) contains a large number of eukaryote-expressed proteins individually printed on a single array slide. 19,394 unique proteins representing ∼75% of the human proteome were evaluated. In total, 14,906 proteins could be assigned to cell physiological processes (**▸Fig. 1A**). The microarray slides were probed with biotinylated long-chain polyphosphates (500 µM, [4]) followed by fluorescence detection (**▸Fig. 1B;** full dataset available at: https://doi.org/10.5281/zenodo.5748254).

**Figure 1.**
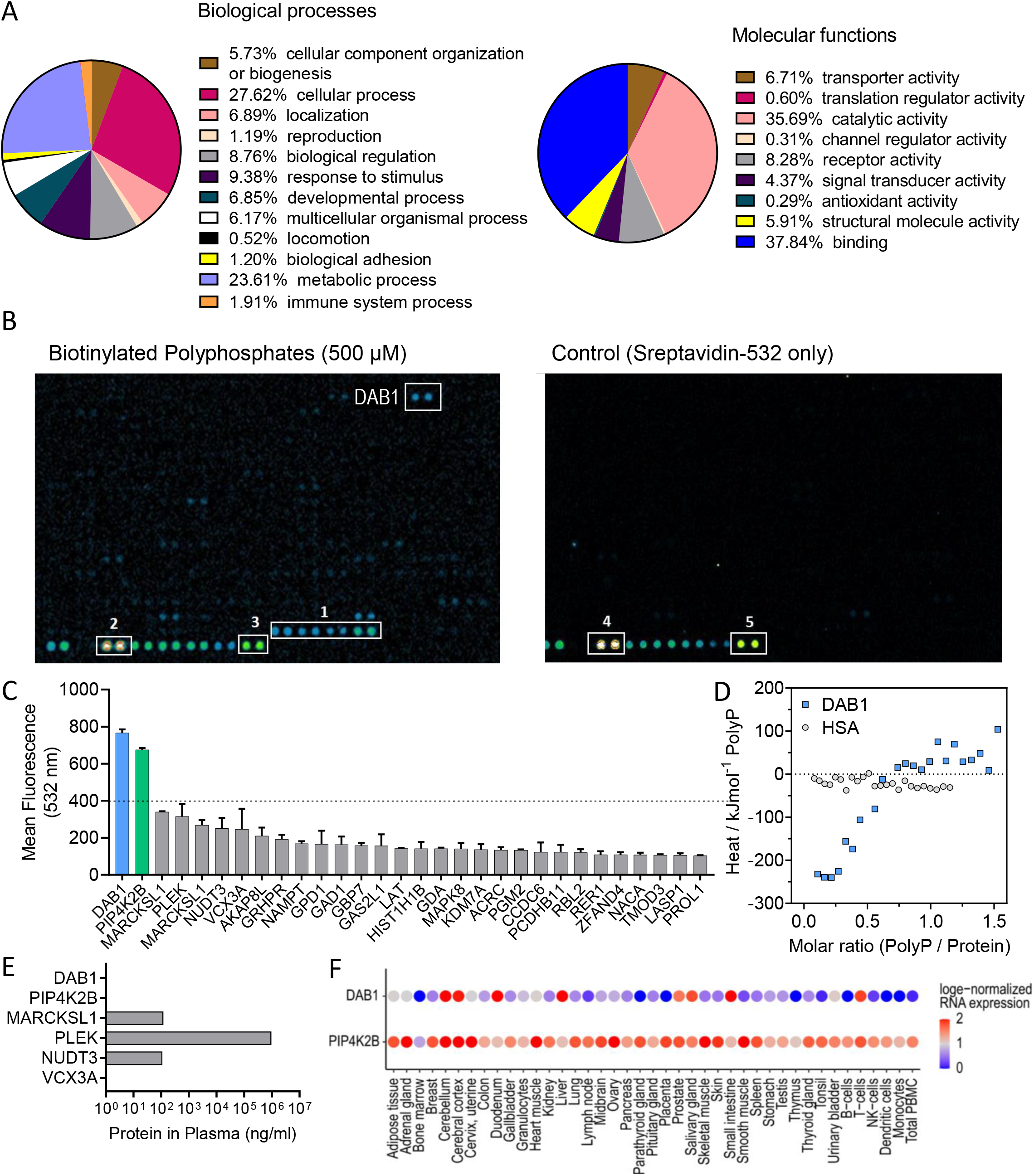
A human proteome microarray for the identification of polyphosphate-binding proteins. (**A**) Overview of the proteins in the HuProt™ human proteome microarray. In total 19,394 human proteins were assessed and 14,906 (multiple mapping possible) were assigned to cell physiological processes using the PANTHER software. (**B**) Representative microarray slides probed with polyphosphates and controls showing the best hit DAB1. Slides were probed with biotinylated long-chain polyphosphates (Pi∼700), followed by a fluorescence detection at 543 nm excitation. The left array shows the subarray with polyphosphates and the right panel depicts a control array. The bottom rows of the subarrays contain control spots; box 1: histones; box 2,4: biotinylated bovine serum albumin; box 3,5: directly fluorophore labelled IgG. No specific protein interactions were observed for the subarray on the control slide (right panel). (**C**) Mean fluorescence intensities of polyphosphate binding signals to the top 30 proteins in the microarray. The horizontal dashed line indicates the assigned threshold. (**D**) Isothermal titration calorimetry measurements of long-chain polyphosphates (20.8 µM in OPCA or PBS buffer) titrated to either recombinant human DAB1 (4 µM in OPCA buffer) or human serum albumin (HSA, negative control, 3 µM in PBS). The integrated heat of a single series of measurements is shown. For calculating molar ratios, a length of 700 phosphate residues per chain was applied. (**E**) Concentrations of selected proteins in human plasma. The data was retrieved from the open-access Peptide Atlas (plasma non-glyco 2017) based on mass spectrometry proteomics. (**F**) Dot plot showing mRNA expression of DAB1 and PIP4K2B in human tissues and immune cells. Consensus normalized expression was retrieved from the Human Protein Atlas (HPA) and is based on combined transcriptome data from the HPA, Genotype-Tissue Expression (GTEx) and Functional Annotation of Mammalian Genomes 5 (FANTOM5) projects. The colors represent the natural log (loge)- normalized RNA expression values.[12]

The best binding partner for polyphosphates was the protein DAB1 (Disabled-1; **▸Fig. 1C**). While DAB1 is a key player of the Reelin-DAB1 signaling pathway, regulating the mammalian brain development by orchestrating cell migration, additional roles in non-neuronal tissues have become more evident in recent years.[5, 6] DAB1 functions as an intracellular adaptor for the canonical binding of the extracellular matrix glycoprotein Reelin to ApoER2 (Apo E receptor 2) and VLDLR (very low-density lipoprotein receptor). Intracellular phosphorylation of DAB1 by Src family Tyrosine kinases (Fyn/Src) facilitates downstream effector recruitment like PI3Ks (phosphatidylinositol 3-kinases), resulting in activation of multiple cellular signaling cascades. DAB1 interacts with PI3Ks in response to Reelin signaling.[7] DAB1 is phosphorylated after binding of activated protein C and PAR activation upstream of PI3Ks.[8] Reelin is also expressed by platelets and it was described that Reelin-platelet interactions can enhance platelet spreading on Fibrinogen.[6, 9] Reelin is important in coagulation, but may also exert functions in T cells and macrophages.[10, 11]

We confirmed the interaction of polyphosphates with human DAB1 using isothermal titration calorimetry (ITC) at 25 °C. As shown in **▸Fig. 1D**, polyphosphates interacted with recombinant DAB1 in an exothermic process with a heat release in the order of magnitude, which is usually observed for protein-protein interactions. However, it was not possible to fit the obtained data due to the low number of titration measurements performed with the limited amount of protein available. We used titration of polyphosphates to human serum albumin (HSA) as a negative control without signs of binding as no significant heat evolvement was detectable (**▸Fig. 1D**). DAB1 is an intracellular protein and not present in human plasma (**▸Fig. 1E**). This suggests that polyphosphates released from platelets, mast cells or bacteria to the extracellular spaces may possibly cross cell membranes to find their intracellular protein targets. In fact, this concept is supported by our recent finding of rapid uptake of fluorescent-labeled polyphosphates into macrophages.[3] DAB1 gene expression was enriched in the brain, the digestive system and in T cells (**▸Fig. 1F**).[12] While sparse information is available on polyphosphates and T cells, polyphosphates are mediators of astroglial signal transmission in mammalian brain [13].

The proteome microarray screen also indicated the binding of polyphosphates to PIP4K2B (phosphatidylinositol-5-phosphate 4-kinase 2B; **▸Fig. 1C**). PIP4K2B is an intracellular protein and absent in human plasma (**▸Fig. 1E**). PIP4K2B is ubiquitously expressed in many tissues (**▸Fig. 1F**). PIP4K2B is a stress-regulated lipid kinase that catalyzes the phosphorylation of phosphatidylinositol-5-phosphate to form phosphatidylinositol-5,4-bisphosphate (PIP2), which is a substrate for the phosphatidylinositol signal transduction pathway. Loss of PIP4K2B is associated with slower tumor growth in p53-deficient mice.[14, 15] Our efforts to study the interactions of polyphosphates and human PIP4K2B using ITC were unsuccessful because the recombinant PIP4K2B was poorly soluble, which is likely a consequence of a lipophilic structure given its localization in membranes.

Several other binding proteins were observed in the microarray with detectable affinity, albeit with lower scores than the threshold recommended by the manufacturer:

MARCKSL1 (Myristoylated alanine-rich C-kinase substrate-like 1, **▸Fig. 1C**) can sequester PIP2 within “lipid rafts” of the cell membrane for participation in later signal transduction events.[16] MARCKSL1 plays a critical role in the regulation of apoptosis and promotes anti-angiogenic effects by reducing VEGF and HIF-1α.[17]

PLEK (Pleckstrin) is highly expressed in blood cells, especially platelets, bone marrow, lymphoid tissue and present in blood plasma (**▸Fig. 1C,E**).[12] In platelets, PLEK is a major substrate of protein kinase C. Upon binding to PIP2 it is involved in the regulation of multiple platelet processes, e.g., aggregation, degranulation, platelet activation and secretion.[18, 19]

The NUDT3 (nucleoside diphosphate linked moiety X-type motif hydrolase-3, **▸Fig. 1C**) gene encodes for the diphosphoinositol polyphosphate phosphohydrolase-1. It belongs to the MutT, or Nudix, protein family and cleaves a β-phosphate from the diphosphate groups in diphosphoinositol pentakisphosphate and bisdiphosphoinositol tetrakisphosphate, suggesting that it may play a role in signal transduction.[20] NUDT3 was identified in a gain-of-function screen as a PI3K inhibitor resistance gene suggesting a role in this pathway.[21] Interestingly, NUDT3 was recently reported to show polyphosphatase activity with a requirement of Zn^2+^ as cofactor.[22]

NAMPT/visfantin (nicotinamide phosphoribosyltransferase) induces endothelial angiogenesis through activation of the PI3K pathway.[23] The three top 10 polyphosphate-binding proteins VCX3A (Variable charge, X-linked 3A; unknown function), GRHPR (glyoxylate reductase / hydroxypyruvate reductase; metabolism) and GPD1 (glycerol-3-phosphate dehydrogenase; metabolism; **▸Fig. 1C**) have no documented connection to the phosphatidylinositol pathway.

In summary, 7 out of the top 10 proteins with highest polyphosphate binding were associated with the PI3K pathway. Interestingly, polyphosphates directly modulate PI3K/Akt signaling [24], although most of the proteins above were not yet known as interaction partners before. On the other hand, our microarray screen could not confirm binding of polyphosphates to their putative receptors, RAGE and P2Y1 [13, 25], which is in line with their non-significant role for mediating polyphosphate effects in macrophages [3]. The number of binding proteins in our screen was lower than expected and provides clear directions for future studies on the molecular targets of polyphosphates.

## Acknowledgements

We thank the team of Cambridge Protein Arrays Ltd., Babraham Research Campus, Cambridge CB22 3AT, UK for work on the HuProt microarray and comments. We thank James H. Morrissey and Stephanie A. Smith for providing biotinylated polyphosphates. We thank Archana Jayaraman and Johannes Platten for assistance with manuscript preparation and Lindsey Stein for secretarial assistance. This work was supported by financial resources obtained from the Federal Ministry of Education and Research (01EO1503 to M.B.), the Deutsche Forschungsgemeinschaft (BO3482/3-3, BO3482/4-1 to M.B.), the National Institutes of Health (R01AI153613, R01HL141513, R01HL139641 to M.B.), a Marie Curie Career Integration Grant of the European Union (Project 334486 to M.B.), and a Clinical Research Fellowship of the European Hematology Association (to M.B.). C.R. was awarded a Fellowship of the Gutenberg Research College at the Johannes Gutenberg-University Mainz. The authors are responsible for the content of this publication.

